# Loss of factor VIII in zebrafish rebalances antithrombin deficiency but has a limited bleeding diathesis

**DOI:** 10.1101/2024.02.28.582609

**Authors:** Catherine E. Richter, Azhwar Raghunath, Megan S. Griffin, Murat Yaman, Valder R. Arruda, Benjamin J. Samelson-Jones, Jordan A. Shavit

## Abstract

Deficiencies in coagulation factor VIII (FVIII, *F8*) result in the bleeding disorder hemophilia A. An emerging novel therapeutic strategy for bleeding disorders is to enhance hemostasis by limiting natural anticoagulants, such as antithrombin (AT3). To study pro/anticoagulant hemostatic balance in an *in vivo* model, we used genome editing to create null alleles for *f8* and von Willebrand factor (*vwf*) in zebrafish, a model organism with a high degree of homology to the mammalian hemostatic system and unique attributes, including external development and optical transparency. *f8* homozygous mutant larvae surprisingly formed normal thrombi when subjected to laser-mediated endothelial injury, had no overt signs of hemorrhage, but had a modest increase in mortality. We have previously shown that *at3^-/-^* larvae develop disseminated intravascular coagulation (DIC), with spontaneous thrombosis and fibrinogen consumption, resulting in bleeding phenotype marked by secondary lack of induced thrombus formation upon endothelial injury. We found that with loss of FVIII (*f8*^-/-^;*at3*^-/-^), larvae no longer developed spontaneous fibrin thrombi and did produce clots in response to endothelial injury. However, homozygous loss of zebrafish Vwf failed to rescue the *at3* DIC phenotype. These studies demonstrate an altered balance of natural anticoagulants that mitigates FVIII deficiency in zebrafish, similar to human clinical pipeline products. The data also suggest that zebrafish FVIII might circulate independently of Vwf. Further study of this unique balance could provide new insights for management of hemophilia A and von Willebrand disease.

**Key Points:**

□ Zebrafish demonstrate a unique balance of natural anticoagulants that mitigates severe FVIII deficiency.
□ Zebrafish FVIII appears to circulate independently of Vwf, which could have implications for management of hemophilia A.

## Introduction

Hemophilia A is an X-linked recessive disease of hemostasis characterized by a deficiency or defect in coagulation factor VIII (FVIII, gene *F8*).^1^ Males are primarily affected, occurring in 1 in 5000 live male births globally.^2^ The cause can be attributed to nonsense, insertion, deletion, splice site, and inversion mutations of the *F8* gene.^1,3^ Individuals with severe hemophilia A exhibit spontaneous bleeding into the soft tissue, joints, and muscles that leads to morbidity and mortality if untreated.^4^ Although prophylaxis with recombinant FVIII is an effective treatment option, some individuals develop neutralizing alloantibodies and experience recurrent bleeding.^5,6^ Novel therapies include FVIII mimetics and modulation of hemostatic balance through targeting natural anticoagulants.^7,8^

Human FVIII is predominantly produced in liver endothelial cells and plays a key role in propagation of the coagulation cascade.^9–11^ The *F8* gene resides in the Xq28 region and has 26 exons.^12^ FVIII has a A1-A2-B-A3-C1-C2 domain structure. However, most circulating FVIII is a heterodimer formed by proteolytic cleavages by the paired basic amino acid cleaving enzyme (PACE) furin within the FVIII B-domain; the heterodimer form of FVIII is comprised of a heavy chain (A1, A2, and B domains) and a light chain (A3, C1 and C2 domains) linked together via a metal ion bridge.^13,14^ During proteolytic activation of FVIII by thrombin, the remaining B domain is removed from the heavy chain. Though it compromises about 40% of the protein, the B-domain is dispensable for the hemostatic function of FVIII. As such, recombinant FVIII variants without the B domain (FVIII-SQ) were developed to facilitate recombinant protein expression, but have also been essential for hemophilia A gene therapy.^15,16^ FVIII-SQ has been specifically engineered to maintain furin processing and heterodimer formation, though subsequent studies have suggested that the former was actually deleterious for FVIII secretion and activity.^13,14^ FVIII-ΔF is a B-domain deleted variant that lacks the furin recognition motif and has decreased heterodimer, but increased expression and clotting activity.^13,14^

Von Willebrand factor (VWF) is a multimeric glycoprotein synthesized in endothelial cells and megakaryocytes.^17^ Secreted VWF enables platelet adhesion at the site of injury. In circulation, stable levels of FVIII are maintained by forming a complex with VWF, which prevents rapid clearance of FVIII.^18,19^ The *VWF* gene is located on chromosome 12 and consists of 52 exons.^20^ VWF undergoes processing and propeptide cleavage by furin, the latter being secreted along with mature VWF.^21^ Von Willebrand disease (VWD) is an autosomal dominant inherited bleeding disorder caused by either deficiency or dysfunction of VWF, which equally affects both males and females.^22^ During injury, VWF enables platelet adhesion to the subendothelium, which is one of the earliest events and an integral part of hemostasis.^23^ Besides acting as a carrier, VWF prevents FVIII from executing its procoagulant functions until required at the site of vascular injury. After thrombin cleavage of FVIII, it dissociates from VWF and forms FVIIIa.^24^

Rodent models of the bleeding disorders associated with loss of FVIII and VWF have been developed and studied in great detail. Hemophilia A mouse models exhibit reduced survival and exhibit excessive bleeding after injury.^25,26^ An *F8*^-/-^ rat model created using genome editing zinc finger nucleases demonstrates spontaneous bleeding including central nervous system hemorrhage, which is the most concerning bleeding in hemophilia A patients.^27,28^ *Vwf*-deficient mice exhibit VWD, including spontaneous hemorrhage, prolonged bleeding times, and reduced FVIII levels and activity.^29^ They also exhibit delayed platelet adhesion with diminished thrombus formation.^29,30^

Zebrafish (*Danio rerio*) have been used extensively to study hemostasis and have demonstrated significant homology with the mammalian coagulation system.^31–34^ They possess several distinct advantages over other models including high fecundity, optical transparency, and external development. New developments in genome editing technology have facilitated quick and robust systems for targeted genetic modification,^35–37^ which has paved the way for targeted mutagenesis in zebrafish to study coagulation disorders.^38^ Here we report targeted ablation of zebrafish *f8* and *vwf* using CRISPR/Cas9-mediated genome editing. We find that zebrafish possess a unique balance within their coagulation system, enabling them to withstand the loss of *f8* without an overt hemorrhagic phenotype yet a modest decrease in their long-term survival, and deletion of *vwf* without apparent phenotype. However, FVIII deficiency is also able to rescue the previously described prothrombotic phenotype of antithrombin (*at3*) deficiency.^39^ This highlights the adaptability of zebrafish in maintaining hemostasis even in the absence of critical procoagulant clotting factors.

## Methods

### Zebrafish strains, maintenance, and survival curves

Zebrafish were raised in accordance with animal care guidelines as approved by the University of Michigan Animal Care and Use Committee. Embryos are defined as 0 to 2 days post fertilization (dpf), larvae 3 to 29 dpf, juvenile 30 to 89 dpf, and adult from 90 dpf.^40^ *f8* and *vwf* mutant zebrafish were created on an ABxTL hybrid background as were the antithrombin (*at3*) mutant and fibrinogen beta chain-enhanced green fluorescent protein (*fgb-egfp*) transgenic lines derived in previous studies.^39,41^ All studies were performed on clutchmates form heterozygous incrosses unless otherwise noted.

### Targeted mutagenesis of the *f8* and *vwf* locus using CRISPR/Cas9 genome editing

The *f8* and *vwf* genes were located in the zebrafish genomic sequence. Target sites of 17-20 nucleotides were identified using ZiFiT (http://zifit.partners.org) as described.^42^ For *f8*, templates for synthetic single guide RNAs (sgRNAs) were constructed with PCR fusion of overlapping oligodeoxynucleotides (IDT, Integrated DNA Technologies, Coralville, IA) which contained the complementary 20 nucleotides of the target site in exon 4, T7 promoter, and sgRNA backbone sequences (Supplementary Table 1), and used for in vitro transcription as described.^43^ Cas9 mRNA was transcribed as previously described.^36^ sgRNAs and Cas9 mRNA at concentrations of 12.5 and 300 ng/µL, respectively were injected into one-cell stage embryos. For *vwf*, crRNAs were designed targeting exons 5, 14, 28, and 51 and purchased (IDT). To produce ribonucleoprotein complexes (RNPs), specific crRNA and tracrRNA were mixed in a 1:1 molar ratio, heated to 95°C for five minutes, cooled to room temperature, and then incubated with Cas9 protein (IDT) at 37°C for 10 minutes. Assembled RNPs were injected into one-cell stage embryos. Injected embryos were raised to adulthood and outcrossed to assess for germline transmission. Deletions were confirmed with Sanger sequencing. *vwf^-^*^/-^ refers to the large deletion generated from exons 5 to 51, except where noted. The exon 14-28 deletion allele is annotated as *vwf*^14–28^^Δ^.

### Genotyping of mutant offspring

Zebrafish larvae or adults were anesthetized in tricaine (0.16 mg/mL, Western Chemical Inc.) or humanely killed in either high dose tricaine (1.6 mg/mL) or by immersion in 4℃ ice water. Tail fin biopsies were obtained as described.^39,44^ Biopsies were incubated in lysis buffer (10 mM Tris-Cl, pH 8.0; 2 mM EDTA, 2% Triton X-100, and 100 µg/mL proteinase K) at 55°C for 2 or more hours, and proteinase K was inactivated at 95°C for five minutes.^39^ PCR was performed on isolated genomic DNA followed by analysis on a QIAxcel Advanced System instrument using capillary electrophoresis (Qiagen). Primer3plus^45^ was used to design primers and oligonucleotides obtained from IDT (Supplementary Table 1).

### Laser-induced endothelial injury in zebrafish larvae

Laser-mediated endothelial injury (MicroPoint Pulsed Laser System, Andor Technology) was performed in the venous system (posterior cardinal vein, PCV) at 3 dpf or arterial system (dorsal aorta, DA) at 5 dpf as described.^46^ The time to occlusion was recorded for up to 2 minutes by an observer blinded to genotype or condition, followed by genotyping.

### Analysis of fibrin deposition

5 dpf *fgb-egfp* larvae (expressing eGFP-labeled fibrinogen^41^) were embedded in low melt agarose, mounted on glass cover slips, and scored for fibrin deposition in the PCV by observers blinded to genotype. Thrombi were semi-quantitatively scored based on the number and size of GFP-labeled fibrin clots, 0: no GFP, 1: less than 5 individual clots, 2: 5-25 individual GFP clots of varying sizes distributed evenly, and 3: continuous large fibrin clots along the vessel. After scoring, larvae were genotyped.

### Gene expression

Using the Zebrafish Embryonic Genotyper (ZEG, Daniolab), cells were extracted from 3 dpf larvae for genotyping. Three biological replicates of each genotype consisting of 1-2 larvae were isolated at 33 dpf, RNA extracted using RNeasy (Qiagen), followed by reverse transcription with Superscript III (Invitrogen). Reverse transcription quantitative real-time polymerase chain reaction (RT-qPCR) was performed on an Applied Biosystems StepOnePlus™ using Fast SYBR (Thermofisher). The relative change in gene expression was analyzed in triplicate using the 2^-ΔΔCT^ method^47^ with *β-actin* as the reference gene.

### Whole mount in situ hybridization (WISH)

Embryos were treated with 30% Danieau/1-phenyl-2-thiourea (PTU) at 6-8 hours post-fertilization (hpf) to inhibit pigmentation. The medium was exchanged every 24 hours until 5 dpf larvae were harvested and fixed in 4% paraformaldehyde (PFA) in phosphate-buffered saline (PBS) overnight at 4°C and stored at -20°C in 100% methanol. Partial *f8* cDNA fragments were amplified to synthesize antisense and sense digoxigenin (DIG)-labeled riboprobes with T7 promoter overhangs primers (Supplementary Table 1). These templates were transcribed *in vitro* using T7 and digoxigenin-RNA labeling mix (Roche). The quality of the synthesized DIG-labeled RNA probes was confirmed by electrophoresis and dot-blot, followed by WISH as described previously^48,49^. In brief, antisense and sense riboprobes were heated to 80°C for 5 minutes and chilled immediately on ice for 5 minutes, incubated overnight in hybridization buffer at 65°C. After hybridization, larvae were washed in PBS with 0.1% Tween 20, re-fixed in 4% PFA, blocked with sheep serum, incubated with anti-digoxigenin-AP fab fragments in blocking buffer, and stained with NBT/BCIP solution. Stained larvae were equilibrated in 70% glycerol and visualized for *f8* expression using a stereomicroscope (Leica) and photographed with an Olympus DP22 2.8-megapixel digital camera.

After WISH, stained larvae were fixed in 4% PFA in PBS for 20 minutes, washed 3x for 5 minutes with PBS, transferred to 20 ml scintillation glass vials containing 30% sucrose in PBS, and incubated at 4°C overnight. 6-10 larvae were placed in molds (Peel-A-Way disposable embedding molds, Polysciences) and sucrose solution removed. Larvae were embedded and frozen in optimal cutting temperature (OCT) compound (Fisher). 10 μm thick cryosections were transferred onto clean glass slides (Fisherbrand Super Frost Plus), dried at 37°C for 1 hour, and stored at 4°C. The dried slides were fixed in 4% PFA, washed 3x with PBS, mounted with permount, and imaged using a Leica DM 5000B microscope.

### Construction of *f8* expression vectors

The pT2AL-*ubi*-*egfp* plasmid was digested with NcoI and ClaI to remove *egfp.*^50^ Zebrafish *f8* cDNA was amplified from total zebrafish cDNA, cloned using CloneJET™ PCR Cloning Kit (ThermoScientific, USA), amplified using primers with homology to pT2AL-*ubi*-*egfp,* followed by HiFi cloning (NEB) to generate pz*f8*. The human B-domain deleted and furin deleted *F8* variants were cloned under control of the SV40 promoter, labeled as ph*F8*-SQ and ph*F8*-ΔF respectively.^13^

### *In vivo* rescue assays

pz*f8* with transposase mRNA, ph*F8*-SQ, and ph*F8*-ΔF were each injected into one-cell embryos generated from *f8*^-/-^;*at3*^-/-^;*fgb*-*egfp* incrosses. pfz*f8* was also injected into embryos generated from *f8*^+/+^;*at3*^+/+^ and *vwf*^14–28^^Δ/Δ^;*at3*^+/-^ incrosses in the *fgb*-*egfp* background. At 5 dpf larvae were scored for fibrin deposition followed by genotyping.

### Statistical Analyses

Graphs, survival curves, and significance testing were generated using Prism (GraphPad Software, California). Time to occlusion (TTO) was analyzed by Mann-Whitney *U* testing, and survival curves compared using log-rank testing (Mantel-Cox). Ordinal scale statistics were performed by comparing experimental groups with the “uninjected” controls based on the fibrin deposition scores. For this purpose, an ordinal library was utilized in deriving cumulative link models. Marginal means were estimated from these models and Dunnett adjustments were applied for multiple comparisons by using *emmeans* library. Analyses were performed in R (v4.2.2).

## Results

### Targeted disruption of *f8* using genome editing nucleases shows no apparent phenotype

To create a model of FVIII deficiency, we used CRISPR-Cas9 genome editing to target exon 4 of the zebrafish *f8* genomic locus (Figure 1A). Injected embryos were raised to adulthood and mated to wild type fish to identify potential F0 founders with a frameshift mutation. A 113 bp deletion (*f8*^-/-^) was confirmed in genomic DNA, and qualitative RT-PCR from cDNA using primers flanking the deletion revealed no product (Figures 1B-D). RT-qPCR data of *f8* expression levels revealed a 90% decrease in the homozygous mutant embryos (Figure 1E). Heterozygotes were incrossed, and genotyping at 3 and 107 dpf revealed a normal Mendelian distribution. Loss of homozygous *f8* mutant fish was not observed until 174 dpf (Figure 1F), but one year there is a small, but statistically significant loss of homozygotes (28%, p=.0015 by log-rank Mantel-Cox) (Figure 1F).

**Figure 1.**
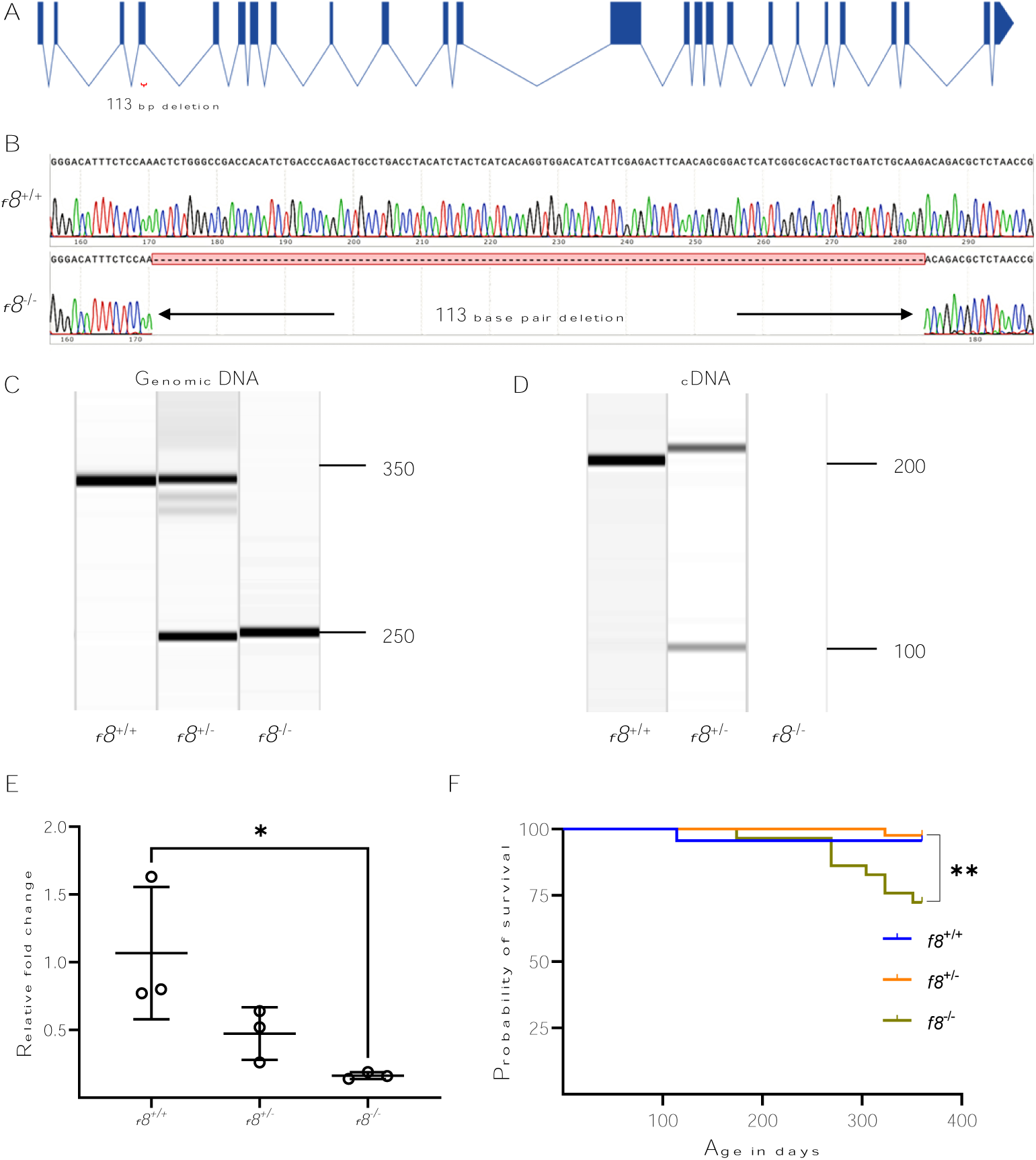
Targeting of the *f8* locus produces a null allele. (A) Structure of the *f8* zebrafish gene and location of the exon 4 deletion. (B) Sequencing of *f8*^-/-^ cDNA demonstrates a 113 bp deletion resulting in a frameshift mutation with a premature stop codon. Amplification of (C) genomic DNA and (D) cDNA from *f8*^+/-^ incrosses. (E) RT-qPCR performed in *f8*^+/-^ incross; *f8*^-/-^ fish show a significant decrease of 90% (± 28%) expression levels compared to *f8*^+/+^. The mean CT values were compared to the reference gene *β-actin* to calculate the relative fold change. Error bars represent standard deviation and statistical significance was determined by a Student’s t test (p = 0.03). (F) Survival curves of zebrafish offspring from an *f8* heterozygous incross starting at 3 months of age shows a statistically significant loss of 25% of homozygotes by one year of age (**p<0.002 by log-rank Mantel-Cox testing), *f8*^+/+^ n=23, *f8*^+/-^ n=42, *f8*^-/-^ n=29.

### Induced clot formation is normal in *f8* mutants but modifies *at3*-deficient coagulopathy

Despite the loss of FVIII and late lethality, there were no grossly observable bleeding in homozygous mutants. Furthermore, there was no statistical difference in the time to occlusion following laser-mediated venous endothelial injury in the PCV between *f8^-/-^* larvae and their heterozygous clutchmates (Figure 2A), with all demonstrating the wild-type response. The lack of an observable bleeding phenotype in FVIII deficient fish led us to wonder if we might uncover a defect when combined with another coagulation factor mutation. Larvae deficient in At3 do not form thrombi in response to endothelial injury, which we previously demonstrated is due to disseminated intravascular coagulation (DIC)-like spontaneous consumptive coagulopathy that consumes fibrinogen.^39^ We hypothesized that a deficiency in FVIII, a procoagulant, could potentially reduce spontaneous clotting events in the *at3^-/-^* mutants and prevent fibrinogen consumption. We crossed *f8* and *at3* double heterozygotes and evaluated them with laser-mediated endothelial injury (Figure 2B). We observed that *f8*^+/+^;*at3*^-/-^ larvae did not occlude within two minutes, consistent with our earlier studies.^39^ However, *f8*^-/-^;*at3*^-/-^ mutants demonstrated normal occlusion indicating rescue of the *at3* DIC phenotype. The time to occlusion for *f8*^+/-^;*at3*^-/-^ mutants showed an intermediate phenotype. These results are consistent with rescue of the *at3*^-/-^ DIC phenotype.

**Figure 2.**
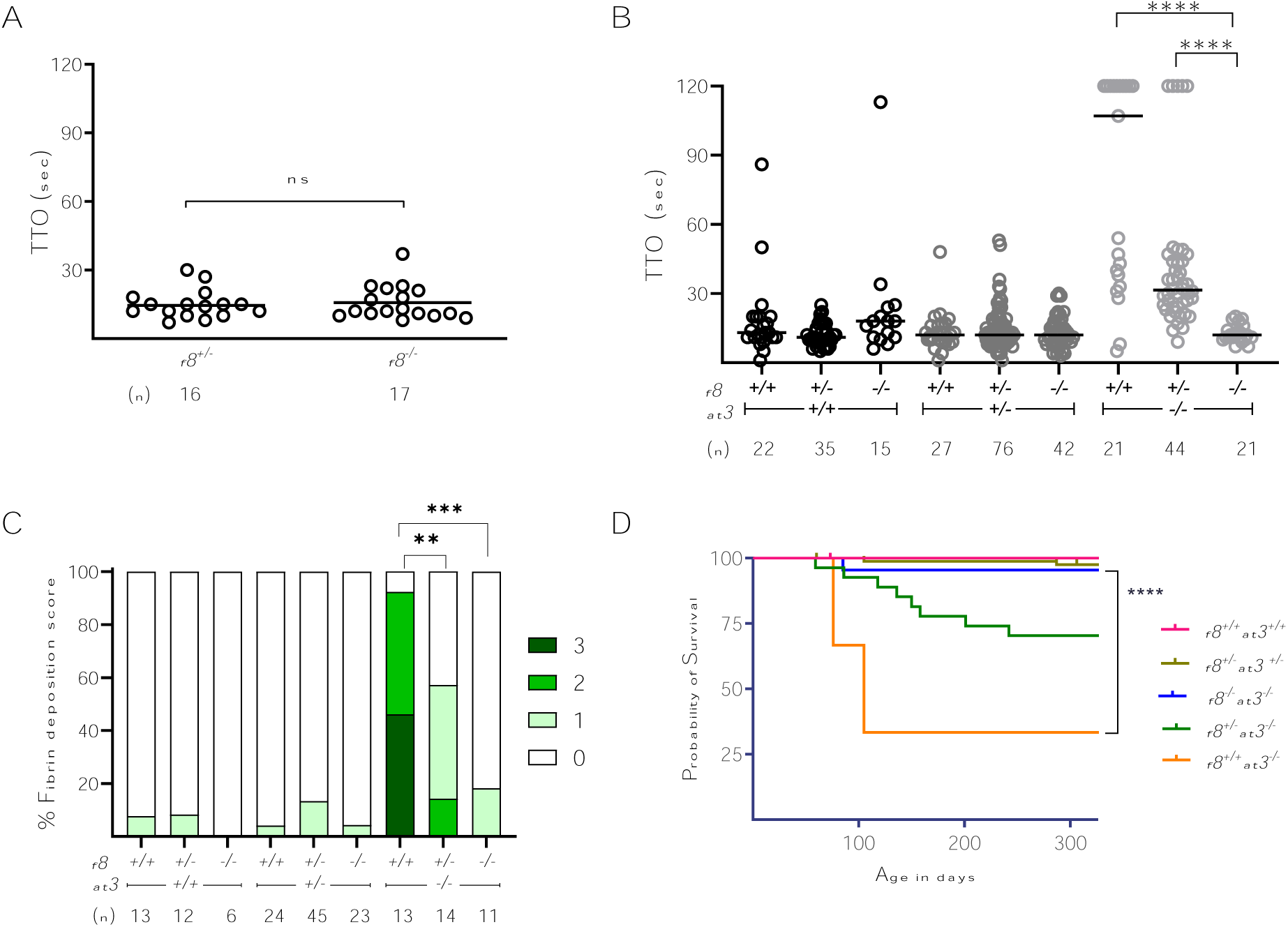
*f8* mutants show an altered hemostatic balance compared to mammals. (A) FVIII deficient zebrafish laser-mediated endothelial injury of the PCV was performed on larvae at 3 dpf. The TTO was not significantly different in *f*8^-/-^ compared to *f*8^+/-^ clutchmates (Mann-Whitney *U* test p > 1). Circles represent individual larvae. Horizontal bars represent the median time to occlusion. (B) Laser-mediated injury on *f8*^+/-^*;at3^+/-^* incrosses reverse the *at3^-/-^* phenotype. In the *at3^-/-^* background, the increased TTO is reversed by mutation of *f8* in a dose-dependent fashion (**** p < 0.001 by Mann-Whitney *U* testing). (C) Fibrin deposition observed in 5 dpf *fgb-egfp* larvae resulting from *f8*^+/-^*;at3^+/-^* incrosses. Bar graph represents the percentage of larvae in each fibrin deposition category: score 0 having no GFP labeled fibrin deposits in the PCV, score 1, less than 5 occurrences, score 2 with 5-25 occurrences, and score of 3 with widespread continuous threads of fibrin in the PCV and/or surrounding regions. Overall statistical significance was determined by Kruskal-Wallis and pairwise comparisons among the *f8*^+/+^;*at3*^-/-^, *f8*^+/-^;*at3*^-/-^, and *f8*^-/-^;*at3*^-/-^ mean fibrin scores by a Wilcoxon rank sum test with a Bonferroni correction (** p < 0.005, *** p<0.0005). (D) Survival curve of zebrafish offspring from *f8^+/-^;at3^+/-^* incrosses shows that loss of *f8* rescues the *at3*^-/-^ lethal phenotype with a statistically significant difference between *f8^+/+^;at3^-/-^* and *f8^-/-^;at3^-/-^* (**** p<0.0001 by log-rank Mantel-Cox testing). *f8^+/+^;at3^+/+^* n=22, *f8^+/-^;at3*^+/-^ n=83*, f8^+/+^;at3^-/-^* n=3*, f8^+/-^;at3^-/-^*, n=27*, f8^-/-^;at3^-/-^*, n=22.

To determine whether this rescue also affected spontaneous PCV thrombosis observed in *at3^-/-^* mutants,^39^ a transgenic EGFP-labeled fibrinogen line (*fgb-gfp*) was bred into the *f8*^+/-^;*at3*^+/-^ mutant background, followed by incrosses. 92% of *f8*^+/+^;*at3*^-/-^ showed elevated levels of fibrin deposition (Figure 2C), consistent with the expected DIC phenotype. Almost none of the *f8*^-/-^;*at3*^-/-^ mutants showed fibrin deposition, and *f8*^+/-^;*at3*^-/-^ mutants once again had an intermediate phenotype. These results are consistent with and parallel our endothelial injury data.

Previously, we have shown that less than 20% of *at3* homozygous mutants survive beyond 7 months of age, the majority dying between 2 and 3 months, in early adulthood.^39^ We tracked offspring from *f8*^+/-^;*at3*^+/-^ incrosses to evaluate long-term survival. This resulted in a significant difference between the survival of *f8*^-/-^;*at3*^-/-^ and *f8*^+/+^;*at3*^-/-^ (Figure 2D), with loss of FVIII rescuing the *at3* lethal phenotype. *f8*^+/-^;*at3*^-/-^ mutants showed partial survival, while *f8*^-/-^;*at3*^-/-^ lived well into adulthood and were indistinguishable from wild-type.

### Expression of *f8* in 5 dpf wild-type zebrafish larvae

We analyzed zebrafish *f8* mRNA expression in 5 dpf wild-type zebrafish larvae by whole-mount in situ hybridization (WISH). *f8* expression was localized to the heart (Figure 3A). Cryosections were performed in the 5 dpf WISH zebrafish larvae. Both the sagittal (Figure 3B) and transverse (Figure 3C) sections revealed *f8* expression in the atrial and ventricular walls of the heart, suggestive of endothelial expression. These results of zebrafish *f8* expression correspond to single-cell (scRNAseq) gene expression data generated from zebrafish embryos and larvae available in three online resources.^51–54^ Collectively, these scRNAseq data indicate expression in the endothelial cells of the heart, consistent with our data. Taken together, we conclude that endothelial cells are the source of *f8* expression in zebrafish larvae, consistent with the source of expression seen in the mouse.^9,10,55^

**Figure 3.**
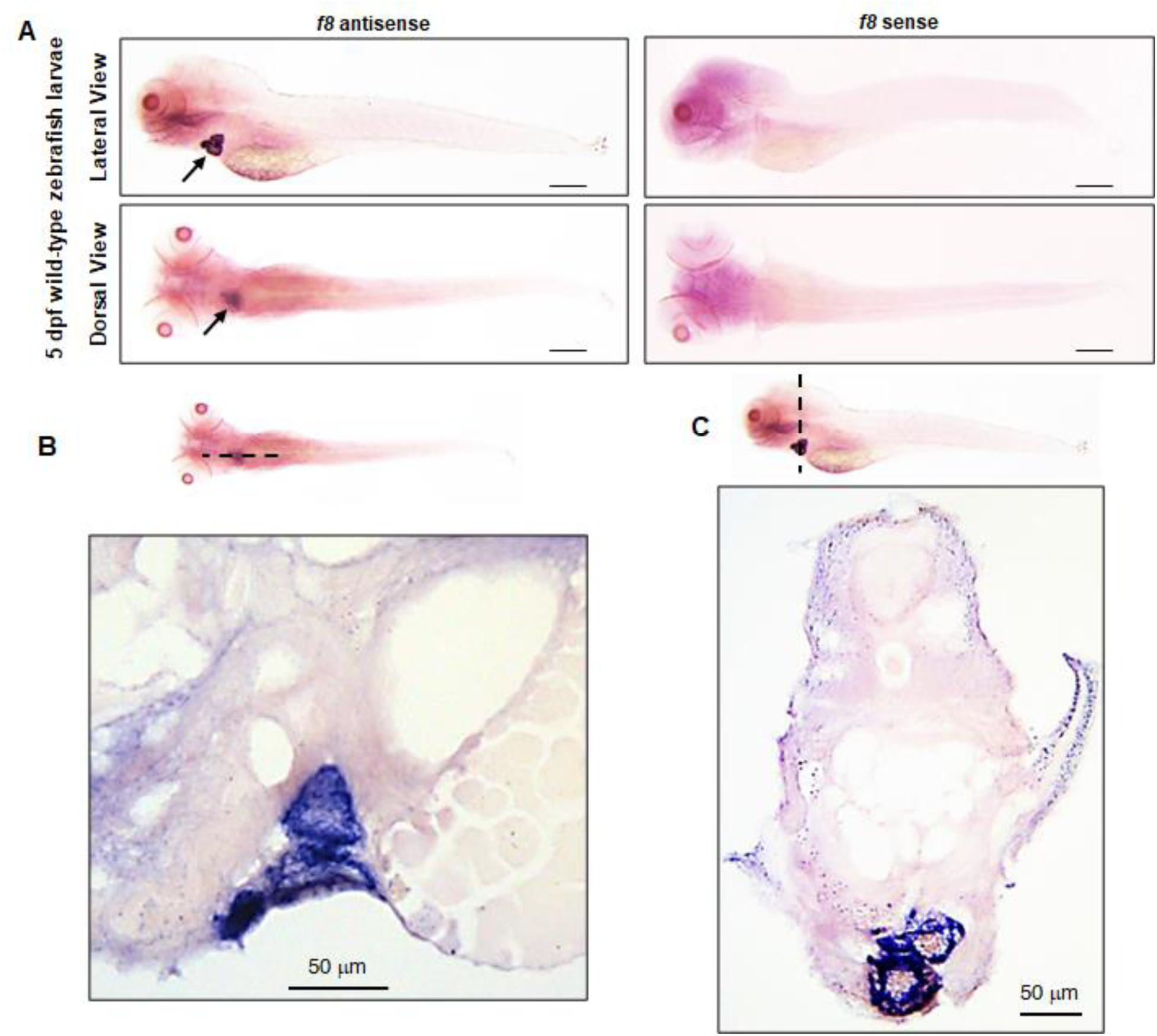
Whole mount *in situ* hybridization (WISH) of *f8* expression in wild-type zebrafish larvae at 5 dpf. (A) *In situ* hybridization with an antisense probe shows *f8* expression in the heart, while a sense controls probe shows no signal. Scale bars, 5 mm. *f8* expression is located in the atrial and ventricular walls of the heart as shown in (B) sagittal and (C) transverse sections (10 µM) of stained larvae. Scale bars, 50 µm.

### Zebrafish and human FVIII restore spontaneous fibrin deposition

We evaluated several forms of FVIII expression in *f8*^-/-^;*at3*^-/-^ mutants through injection of plasmids encoding the cDNAs for zebrafish *f8* (pz*f8*), human B-domain deleted *F8* (ph*F8*-SQ), and human B-domain deleted high-expressing *F8* (ph*F8*-ΔF) variant. Injection with either pz*f8*, ph*F8*-SQ, or ph*F8*-ΔF into *f8*^-/-^;*at3*^-/-^ mutant embryos significantly restored thrombosis formation to roughly equivalent levels as the *f8*^+/+^;*at3*^-/-^ mutants (Figure 4A, 4B, and 4C respectively). Zebrafish *f8* (pz*f8*) appeared to be more efficient than the human *F8* cDNAs, which were roughly equivalent to one another, though there was a non-significant trend of increased activity for ph*F8*-ΔF. pz*f8* injected into wild-type larvae similarly exhibited thrombus formation similar to the *f8*^+/+^;*at3*^-/-^ mutant phenotype (Figure 4D).

**Figure 4.**
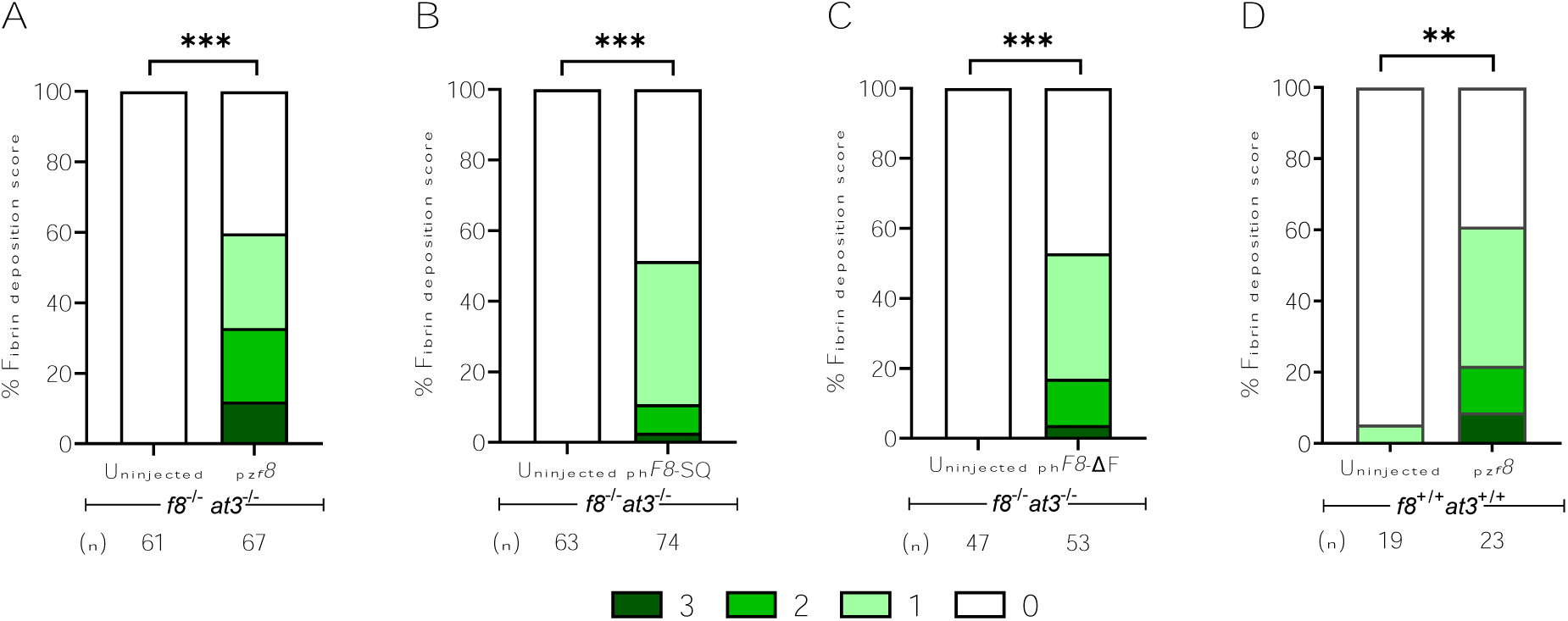
Overexpression of FVIII results in thrombosis. Either *f8^+/+^*;*at3^+/+^*or *f8^-/-^*;*at3^-/-^* zebrafish, both in the *fgb-gfp* background, were incrossed and one-cell stage embryos injected with pz*f8*, ph*F8*-SQ, or ph*F8*-ΔF and evaluated for spontaneous thrombi formation via fibrin deposition scoring at 5 dpf. pz*f8* injected into (A) *f8^-/-^*;*at3^-/-^*and (D) *f8^+/+^;at3^+/+^* zebrafish larvae exhibited statistically significant spontaneous thrombus formation. (B) ph*F8*-SQ and (C) ph*F8*-ΔF injected into *f8*^-/-^;*at3*^-/-^ zebrafish larvae demonstrated statistically significant levels of spontaneous thrombi. Statistical significance was determined by ordinal logistic regression followed by Dunnett’s adjustments (*** p<0.001; ** p<0.01).

### Laser-induced endothelial injury is normal in Vwf-deficient larvae

We used genome editing to generate two mutants in the *vwf* zebrafish gene; a partial deletion (exon 14-28, *vwf*^14–28^^Δ^) and a nearly whole locus deletion (exon 5-51, *vwf*^-^) (Figure 5A, B, C). When subjected to venous endothelial injury at 3 dpf, or arterial endothelial injury at 5 dpf, both full and partial deletion mutant *vwf^-/-^* larvae did not show a difference in the time to occlusion of the PCV (Figure 5D). No mortality was observed in *vwf*^+/-^ and *vwf*^-/-^ clutchmates tracked to eight months of age (data not shown).

**Figure 5.**
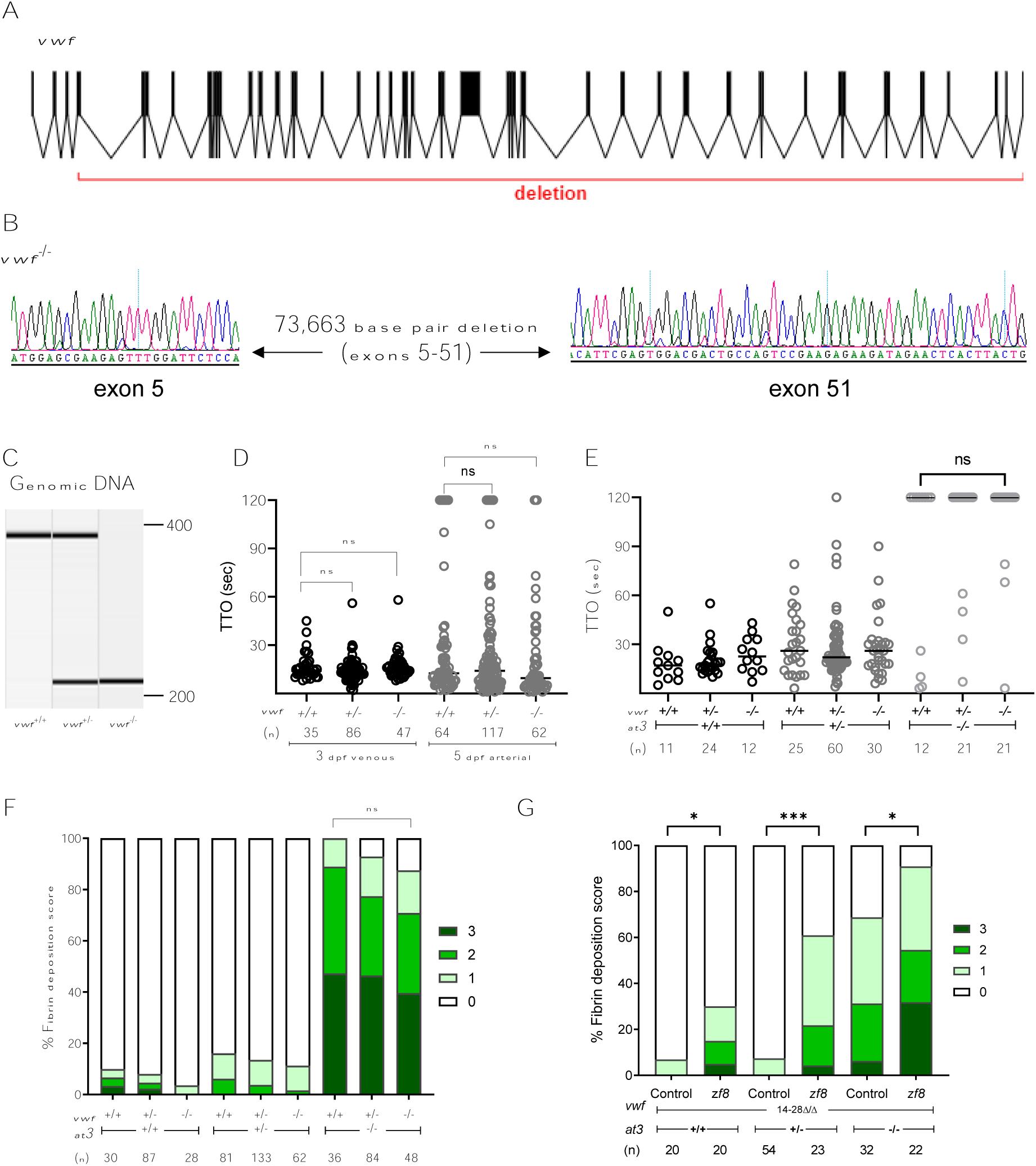
Genome engineering of a large deletion in *vwf*. (A) The structure of the *vwf* gene in zebrafish, with deletion indicated. (B) Sequencing of *vwf*^-/-^ fish shows a large deletion (73.7 kb) between exons 5 and 51. (C) Genomic DNA amplification of *vwf* mutants. (D) Laser-mediated endothelial injury was performed on larvae at 3 dpf (venous) and 5 dpf (arterial). The time to occlusion was not significantly different among *vwf* mutant clutchmates (Mann-Whitney *U*, p=0.5 for venous and p=0.04 for arterial, only values <0.03 are considered significant after Bonferroni correction). Circles represent individual larvae. Horizontal bars represent the median time to occlusion. (E) Laser-mediated injury at 3 dpf from an incross of *vwf*^+/-^;*at3*^+/-^ shows no change in the *at3*^-/-^ DIC bleeding phenotype with loss of Vwf (Mann-Whitney *U*, p=0.12). (F) Spontaneous fibrin deposition observed in 5 dpf larvae from *vwf*^+/-^;*at3*<u>^+/-^</u> incrosses. Bar graph represents percentage of larvae in each category: score 0 having no GFP labeled fibrin deposits in the PCV, score 1, less than 5 occurrences, score 2 with 5-25 occurrences, and score 3 with widespread continuous threads of fibrin in the PCV and/or surrounding regions. Statistical significance was determined by Kruskal-Wallis testing and pairwise comparisons among *vwf*^+/+^;*at3*^-/-^, *vwf*^+/-^;*at3*^-/-^, and *vwf*^-/-^;*at3*^-/-^ (p=0.3). Larvae were confirmed to express the *fgb-egfp* transgene in the liver prior to scoring. (G) p*zf8* injected into one-cell stage embryos collected from *vwf*^14–28^^Δ/Δ^;*at3*^+/-^ incrosses and evaluated for spontaneous thrombi formation via fibrin deposition scoring at 5 dpf. p*zf8* injected zebrafish larvae exhibited significant fibrin deposition compared to uninjected controls in all genotypes. Statistical significance was determined by ordinal logistic regression followed by Dunnett’s adjustments (*** p<0.001; * p<0.05).

Mammalian VWF is known to carry FVIII in circulation and given the conservation of sequence between fish and mammals we suspected that *vwf* mutants would have decreased levels of FVIII. Therefore, *vwf;at3* double mutants should also demonstrate mitigation of the *at3* DIC phenotype, as was the case for *f8;at3* mutants. We crossed both full and partial *vwf* deletions into the *at3* mutant background. Using 3 dpf larvae from a double heterozygous incross we performed venous endothelial injury. Unlike heterozygous and homozygous loss of FVIII, even complete Vwf deficiency did not rescue the *at3*^-/-^ mutant DIC bleeding phenotype (Figure 5E). We also looked for a change in spontaneous thrombus formation in the *vwf*^-^*;at3* double mutants. When combined with *at3* mutants, Vwf deficient larvae showed a slight reduction in fibrin thrombi, but this trend was not statistically significant. (Figure 5F). To determine whether Vwf is required for FVIII in zebrafish circulation, we injected pz*f8* into *vwf*^14–28^^Δ/Δ^;*at3* mutants. We found that injected larvae were still able to increase the level of thrombus formation in the absence of Vwf, regardless of *at3* genotype (Figure 5G).

## Discussion

This study reports the targeted mutagenesis of *f8* and *vwf* in zebrafish. The *f8* mutation is a large exonic deletion causing a frameshift and premature stop codon. The residual truncated transcript is minimally detected, suggesting nonsense mediated decay.^56^ Due to this frameshift mutation and the absence of alternative splice products, it is highly likely that any protein translated from this mutant *f8* gene would be nonfunctional, and therefore we conclude that this is a null allele. We have previously shown severe lethal bleeding defects with loss of extrinsic and common pathway knockouts,^39,43,57–59^ consistent with the analogous mouse mutants. However, while the zebrafish mutants survive into early adulthood before succumbing to the coagulopathy, mouse knockouts present with embryonic or neonatal lethal hemorrhage.

In the case of *f8* mutants, the long-term survival shows similarities with the mouse model. Both have a shortened lifespan attributed to hemorrhagic events.^26^ However, like the other coagulation factor knockouts, the severity was greater in mice, with ∼40% overall survival at one year^26^ versus ∼75% at the same timepoint in our study. Concordantly, while FVIII-deficient mice exhibit impaired clot formation when challenged, we were surprised to find that zebrafish larvae demonstrated no visible hemorrhage and normal hemostasis when challenged with endothelial injury. This may be attributed to a highly tissue factor driven response to endothelial injury that obviates the need for the intrinsic pathway.^57^

In our previous studies, we demonstrated that *at3*^-/-^ zebrafish develop spontaneous thrombosis and uncontrolled DIC during the early stages of development, followed by intracardiac thrombosis and lethality in early adulthood.^39^ We found that there was consumption of fibrinogen, presumably due to unregulated thrombin activity. Here we observed that loss of FVIII counterbalanced the presumed elevated thrombin activity in *at3*^-/-^ mutants, thus rescuing the consumptive coagulopathy, preventing spontaneous thrombosis, and rescuing adults from lethality. This observation aligns with findings in FVIII-deficient mice, where diminished levels of antithrombin were associated with reduced bleeding and increased thrombin generation^60^ as well as a case report where the bleeding phenotype of severe hemophilia A (FVIII activity <1% normal) was ameliorated by AT3 deficiency.^61^ These observations are also consistent with AT3 knockdown, a rebalancing therapy in clinical development for patients with hemophilia.^7^

In mammals, FVIII is primarily synthesized in liver endothelial cells.^9,10^ WISH revealed evidence that expression of *f8* in zebrafish is from the endothelial lining of the heart, and this is confirmed by publicly available scRNAseq data.^51–55,62^ Additionally, single-cell transcriptomic profiling of vascular endothelial cells identified *f8* transcripts among the top 30 differentially expressed genes within the endocardial subcluster.^63^ These data reveal conservation of endothelial cells as the site for FVIII production, although the site has diverged suggesting the possibility of unique adaptations and regulatory mechanisms in fish. We also find the critical role of elevated FVIII levels in thrombosis, contributing to the evidence supporting FVIII as a robust risk factor, aligning with epidemiological data and reinforcing the clinical relevance of our observations.^64,65,66^ This underscores the reproducibility and significance of elevated FVIII levels in triggering thrombotic events across species.

Approved adeno-associated viral (AAV) vectors for hemophilia A gene therapy use the B-domain-deleted variant FVIII-SQ.^16,67^ Pre-clinical mouse and canine studies previously demonstrated a 2-3 fold increase in FVIII levels after AAV gene therapy with h*F8*-ΔF compared to h*F8*-SQ.^13,14^ However, we only observed a small non-significant increase in clot formation with h*F8*-ΔF compared to h*F8*-SQ. This discrepancy may be due to species-specific factors or an experimental system that is insufficiently sensitive to distinguish such small differences. Nevertheless, these findings highlight the overall conservation of FVIII functionality and emphasize the potential for evaluating human functional variants through *in vivo* rescue experiments in zebrafish, as we have done for other coagulation factors.^39,43,58,70^ Bioengineering FVIII variants is one approach to address current limitations of hemophilia A gene therapy.^16^

*Vwf* mouse^29,71^ and rat^72^ knockouts displayed prolonged bleeding after injury, a characteristic feature of patients with VWD. We find that loss of zebrafish Vwf did not affect hemostasis in response to venous and arterial endothelial injury. This is in contrast to previous zebrafish *vwf* knockout studies that reported increased time to occlusion analyzed using laser-mediated and ferric chloride-induced endothelial injury.^73,74^ This could potentially be attributed to the small insertions generated in those studies (7 and 55 bp), while we deleted nearly the entire *vwf* gene. This difference in phenotype might also arise from the nature of *vwf* mutations, with ours representing a clear loss of function, equivalent to type 3 VWD.^19^ The others could possibly have generated dominant negative proteins, mimicking type 1 or 2 VWD subtypes.^75^ Additionally, the lower blood pressure typically observed in zebrafish^76^ compared to mammals may result in altered shear stresses in the vasculature, potentially mitigating the bleeding tendency associated with Vwf deficiency.

The Vwf-deficient *at3* mutants yielded intriguing observations, with absence of rescue of the DIC bleeding phenotype characteristic of *at3* mutants and the lack of any significant alteration in thrombus formation. We had hypothesized that the complete absence of zebrafish Vwf, if it served similar functions to its mammalian counterpart, would result in a severe deficiency of FVIII. Such a deficiency would have logically been expected to reduce thrombosis in *at3* mutant zebrafish, potentially to a similar degree as observed in *f8^+/-^;at3^-/-^* mutants. These findings lead us to speculate that within the zebrafish hemostatic system Vwf might not play a critical role in the carriage and stabilization of FVIII. This raises the intriguing possibility that zebrafish might possess distinct mechanisms for FVIII stabilization or transport. If these mechanism(s) could be applied to humans, they could lead to an enhanced FVIII replacement product for hemophilia A.

In conclusion, our study highlights the remarkable balance in the zebrafish coagulation system, allowing it to tolerate the absence of FVIII and Vwf to a much greater extent than mammals. In contrast to the observed phenotypes in mammals, the loss of Vwf in zebrafish does not appear to manifest with FVIII deficiency. This discrepancy suggests the possibility that zebrafish FVIII has evolved to be stable in circulation without Vwf, and perhaps FVIII-VWF association may be a specific mammalian adaptation. Exploring this distinctive *in vivo* balance holds the potential to uncover new perspectives for managing hemophilia A and VWD.

## Acknowledgements

This work was supported by the National Hemophilia Foundation Judith Graham Pool Postdoctoral Fellowship Award (A.R.) and National Institutes of Health (NIH) grant R35 HL150784 (J.A.S.). B.J.S.-J. was supported by NIH K08 HL140078. J.A.S. is the Henry and Mala Dorfman Family Professor of Pediatric Hematology/Oncology.

## Authorship contributions

C.E.R. and A.R. designed and conducted research, analyzed data, and wrote the manuscript. M.S.G. designed experiments, performed research, and analyzed data. V.R.A. designed research but passed away prior to manuscript preparation. B.J.S.-J. designed research, analyzed data, and edited the manuscript. M.Y. performed statistical analysis. J.A.S. designed research, supervised experiments, conducted research, analyzed data, and wrote the manuscript. All authors reviewed the manuscript.

