## Supplementary Table 1 for "Loss of factor VIII in zebrafish rebalances antithrombin deficiency but has a limited bleeding diathesis"

**Supplementary Table 1**. Oligonucleotide sequences

| Primer | Oligonucleotide Sequences (5'-3') | Description |
|---|---|---|
| 1 | GGGACATTTCTCCAAACTCT | 5' *f8* gRNA amplification primer |
| 2 | GACAGCAACATCCTGAGAGTAAAG | 5' *f8* genotyping |
| 3 | CTGCTTATTTTCCCCGCTGT | 3' *f8* genotyping |
| 4 | CCTTGAATGCCAGCCCATG | 5'cDNA and qPCR *f8* amplification |
| 5 | CGGAGTACACCGATTCCACT | 3' cDNA and qPCR *f8* amplification |
| 6 | **ATTGTAATACGACTCACTATAGGG** TCTCGGTCCTGATACGCTCT | *f8* antisense RNA riboprobe primer (bold represents T7 promoter sequence) |
| 7 | GAAACACAAGCAGGTGCTGA | *f8* antisense RNA riboprobe primer |
| 8 | **ATTGTAATACGACTCACTATAGGG** AAACACAAGCAGGTGCTGA | *f8* sense RNA riboprobe primer (bold represents T7 promoter sequence) |
| 9 | TCTCGGTCCTGATACGCTCT | *f8* sense RNA riboprobe |
|  | GGGACATTTCTCCAAACTCT | *f8* sgRNA exon 4 |
| 10 | GGAGACGACTTCTTAACAGT | *vwf* sgRNA exon 14 |
| 11 | CGCGAGATGCGGTACACCGG | *vwf* sgRNA exon 28 |
| 12 | TCCGTATAGATCCCTTACAT | *vwf* sgRNA exon 5 |
| 13 | GAACTACATTCGAGTGGACG | *vwf* sgRNA exon 51 |
| 14 | GCAGGGTATGAACTGGGCTA | 5' *vwf* genotyping exon 5 |
| 15 | CTGCCACTTTTCCAGTCAAGC | 3' *vwf* genotyping exon 5 |
| 16 | GTATTTGCAGCTAAAACCAGCC | 5' *vwf* genotyping 14 |
| 17 | TTCCTCGCTCTGTTAATGTTGA | 3' *vwf* genotyping exon 14 |
| 18 | CACTGGCACCAACCACCGTC | 5' *vwf* genotyping exon 28 |
| 19 | TGGCTTCATCCATGAATGCG | 3' *vwf* genotyping exon 28 |
| 20 | GGGACGCGTGAACTGGTTAT | 5' *vwf* genotyping intron 50 |
| 21 | ACACAGACTTGCTGCCACAC | 3' *vwf* genotyping exon 52 |
| 22 | CCAAGGCCAACAGGGAAAAG | *β-actin* qPCR 5' |
| 23 | GAGGCATACAGGGACAGCAC | *β-actin* qPCR 3' |
